# Complex carbohydrate utilization by gut bacteria modulates host food preference

**DOI:** 10.1101/2024.02.13.580152

**Authors:** Kristie B. Yu, Celine Son, Anisha Chandra, Jorge Paramo, Anna Novoselov, Ezgi Özcan, Sabeen A. Kazmi, Gregory R. Lum, Arlene Lopez-Romero, Jonathan B. Lynch, Elaine Y. Hsiao

## Abstract

The gut microbiota interacts directly with dietary nutrients and has the ability to modify host feeding behavior, but the underlying mechanisms remain poorly understood. Select gut bacteria digest complex carbohydrates that are non-digestible by the host and liberate metabolites that serve as additional energy sources and pleiotropic signaling molecules. Here we use a gnotobiotic mouse model to examine how differential fructose polysaccharide metabolism by commensal gut bacteria influences host preference for diets containing these carbohydrates. *Bacteroides thetaiotaomicron* and *Bacteroides ovatus* selectively ferment fructans with different glycosidic linkages: *B. thetaiotaomicron* ferments levan with β2-6 linkages, whereas *B. ovatus* ferments inulin with β2-1 linkages. Since inulin and levan are both fructose polymers, inulin and levan diet have similar perceptual salience to mice. We find that mice colonized with *B. thetaiotaomicron* prefer the non-fermentable inulin diet, while mice colonized with *B. ovatus* prefer the non-fermentable levan diet. Knockout of bacterial fructan utilization genes abrogates this preference, whereas swapping the fermentation ability of *B. thetaiotaomicron* to inulin confers host preference for the levan diet. Bacterial fructan fermentation and host behavioral preference for the non-fermentable fructan are associated with increased neuronal activation in the arcuate nucleus of the hypothalamus, a key brain region for appetite regulation. These results reveal that selective nutrient metabolism by gut bacteria contributes to host associative learning of dietary preference, and further informs fundamental understanding of the biological determinants of food choice.

## Introduction

Eating behavior that supports intake of a well-balanced diet is essential for health. The gut microbiota interacts directly with dietary nutrients to regulate host metabolism and energy storage from food^1,2^. Particular microbes specialize in metabolizing different types of nutrients to support their cellular respiration and to ultimately survive in a competitive gut microbial ecosystem^3^. This has led some to hypothesize that gut microbes may manipulate host feeding behavior to promote their own fitness^4^.

Growing evidence supports a role for the gut microbiota in modulating host macronutrient preference. In fruit flies deprived of essential amino acids, gut bacteria suppressed host protein appetite by producing lactate-derived metabolites or amino acids that are detected by gut expression of the neuropeptide CNMamide^5,6^. In mice, colonization with microbiota from wild rodents with different foraging strategies conferred differential food preferences, where those colonized with herbivore-associated microbiota preferred diets with a higher protein to carbohydrate ratio diet than those colonized with omnivore- and carnivore-associated microbiota^7^. Similarly, transplanting microbiota from obese donor mice into lean recipient mice lowered preference for high fat diets, which was correlated with deficits in dopamine signaling in the nucleus accumbens^8,9^. Conversely, treating mice with antibiotics to deplete gut bacteria led them to overconsume palatable high-sucrose diets^10^. These studies provide foundational proof-of-concept that alterations in the microbiome can influence host feeding behavior when comparing diets that vary in macronutrient composition, but whether selective nutrient utilization by gut bacteria impacts host dietary preference remains unclear.

Particular bacterial members of the gut microbiota metabolize complex carbohydrates that are non-digestible by the host^1,2,11^, generating short chain fatty acids (SCFAs) that can reduce host appetite through gut-brain pathways^12–15^. Elevated fecal SCFA levels were correlated with higher subjective hunger ratings and increases in fiber-fermenting bacteria in humans, suggesting that a link between dietary fiber intake, SCFA metabolism, and eating behavior^16^. High colonic levels of the SCFA propionate was associated with reduced activity in the nucleus accumbens during food picture evaluation, decreased subjective appeal of high-energy food pictures, and reduced energy intake during an *ad libitum* meal^17^. Similarly, prebiotic supplementation with inulin in overweight adults decreased reward-related brain activation during food decision-making^18^. Despite these correlations from human studies, whether there are causal relationships between microbial fermentation of complex carbohydrates and host food preference remains unclear.

To investigate this, we focus on species from the genus *Bacteroides*, prevalent human-derived gut bacteria which are critical for digesting complex carbohydrates via genetically-encoded polysaccharide utilization loci (PUL)^19^ to produce the SCFAs acetate and propionate^12^. *Bacteroides thetaiotaomicron* and *Bacteroides ovatus* express PUL variants that restrict their metabolism to fructose-containing polysaccharides called fructans with different glycosidic linkages: *B. thetaiotaomicron* ferments levan-type fructans with β2-6 linkages, whereas *B. ovatus* ferments inulin-type fructans with β2-1 linkages^20,21^. In this study, we use these two phylogenetically related and genetically tractable bacteria in a gnotobiotic mouse model to dissect mechanisms by which bacterial utilization of complex carbohydrates impacts host preference for diets that differ only in carbohydrate type. We further manipulate bacterial PULs to examine causal effects of bacterial fructan utilization on host fructan preference and neuronal activation in the arcuate nucleus of the hypothalamus. Results from this study reveal that selective fructan utilization by gut bacteria shapes host dietary preference and advance mechanistic understanding of how the gut microbiota impacts host feeding behavior.

## Results

### Colonization with fructan-utilizing bacteria promotes host preference for non-fermentable fructan diet

The gut microbiome has the capacity to modify host appetite and preference for diets that vary in macronutrient composition^5–10,13,14^. To determine whether differential fructan utilization by gut bacteria alters host preference for diets that vary only in fructan source, we began by selecting *B. ovatus* and *B. thetaiotaomicron* based on their reported selectivity in degrading inulin and levan^20^, respectively, which are fructans that differ only in their glycosidic linkages (**Fig. 1a**). We generated custom mouse diets containing either 10% inulin or levan purified for dietary formulation (“dietary levan”, see Methods) as the only carbohydrate source (**Table S1**). As expected, *B. thetaiotaomicron* grew in minimal media containing 0.5% of either pure levan or dietary levan, but not inulin as its sole carbohydrate source. *B. ovatus* grew in minimal media containing inulin, but not pure levan (**Fig. 1b**). It also exhibited partial growth on dietary levan, suggesting the presence of residual mono- or oligosaccharides generated from the bulk purification process (**Fig. 1b**). Consistent with this, *B. thetaiotaomicron* grew better in minimal media containing 5% levan diet (LD) than inulin diet (ID), while *B. ovatus* exhibited only a slight growth advantage in minimal media containing ID than LD (**Fig. 1c**, **Extended Data Fig. 1a**). While keeping this caveat in mind, we refer in subsequent sections to LD as a fermentable diet and ID as a non-fermentable diet to *B. thetaiotaomicron*, and LD as a non-fermentable diet and ID as a fermentable diet to *B. ovatus*. We confirmed *in vitro* that *B. thetaiotaomicron* produced more acetate on LD than ID, while *B. ovatus* produced more acetate on ID than LD (**Fig. 1d**).

**Figure 1:**
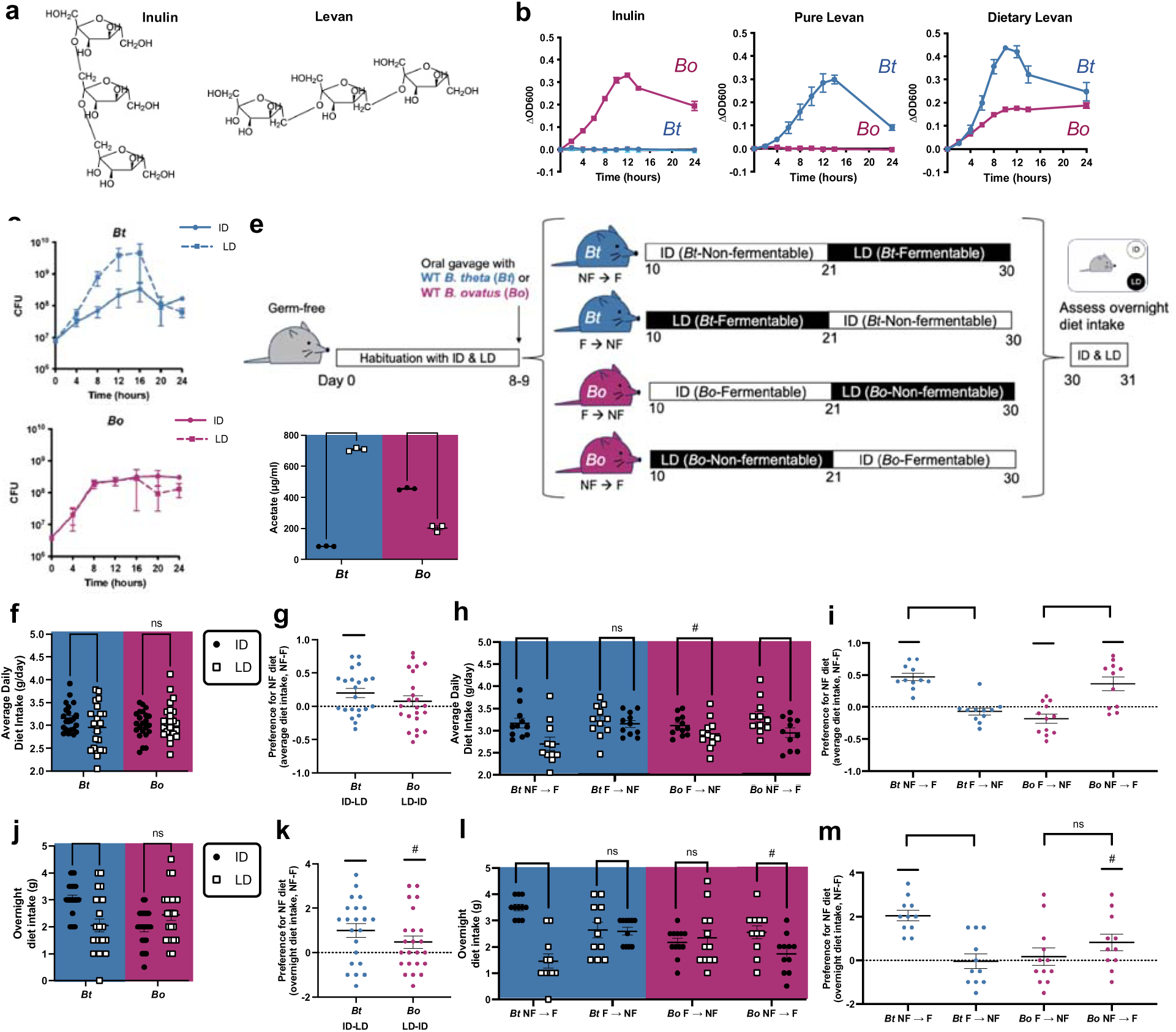
Colonization with differential fructan-utilizing bacteria increases host preference for diets containing the non-fermentable fructan. **a**) Fructan chemical structures. Inulin has *β*2-1 linkages between fructose subunits, while levan has *β*2-6 linkages between fructose subunits. **b**) Growth curves of *B. thetaiotaomicron (Bt)* and *B. ovatus (Bo)* in minimal media containing 0.5% inulin, pure levan, or dietary levan. Each dot represents 3 biological replicates. OD600 = optical density at 600 nm. **c**) Growth curves of *Bt* and *Bo* in minimal media containing 5% inulin diet (ID) or levan diet (LD). Each dot represents 3 biological replicates. CFU = colony forming unit. **d**) Acetate produced in supernatants of bacteria grown for 24 hours in minimal media containing 5%ID or LD. (n = 3 each). **e**) Experimental schematic for sequential feeding post-colonization. Germ-free mice were habituated with both diets for 7 days before being colonized with *Bt* or *Bo*. The mice ate one diet *ad libitum* for 11 days and then the other diet for 10 days. The mice were then provided both diets in their cage overnight. **f**) Average daily diet intake post-colonization for mice colonized with *Bt* and *Bo*. **g**) Preference score for the non-fermentable (NF) fructan in average daily diet intake post-colonization for mice colonized with *Bt* (ID-LD) and *Bo* (LD-ID). **h**) Average daily diet intake post-colonization for mice colonized with *Bt* and *Bo* by diet order. **i)** Preference score for the NF fructan in average daily diet intake post-colonization for mice colonized with *Bt* (ID-LD) and *Bo* (LD-ID) by diet order. **j**) Overnight diet intake from days 30-31 for mice colonized with *Bt* and *Bo*. **k**) Preference score for the NF fructan in overnight diet intake for mice colonized with *Bt* (ID-LD) and *Bo* (LD-ID). **l)** Overnight diet intake from days 30-31 for mice colonized with *Bt* and *Bo* by diet order. **m**) Preference score for the NF fructan in overnight diet intake for mice colonized with *Bt* (ID-LD) and *Bo* (LD-ID) by diet order. Data from f-m are combined from three independent experiments. For f, g, j, and k, n = 22 for *Bt*, 23 for *Bo*. For h, i, l, and m, n = 11 for *Bt* NF ➔ F, 11 for *Bt* F ➔ NF, 12 for *Bo* F ➔ NF, 11 for *Bo* NF ➔ F. For d, f, h, j, and l, 2-way ANOVA with matched column measures, comparing means across rows, and Sidak’s corrections were performed. For g, i, k, and m, 90 and 95% confidence intervals were computed, and significance was determined if the interval did not contain zero. Unpaired parametric t-tests were also performed between groups. # = p-value < 0.10; * = p-value < 0.05; ** = p-value < 0.01; *** = p-value < 0.001; **** = p-value < 0.0001

We hypothesized that differential bacterial fructan utilization would alter host preference for fructan diets. To test this, we monocolonized mice with wild-type *B. thetaiotaomicron* or *B. ovatus*, fed them ID or LD in sequence, and measured their food consumption (**Fig. 1e**). To minimize potential confounds of diet novelty, mice were habituated to both ID and LD for 7 days prior to colonization, and diet order was counterbalanced in each cohort. Throughout the experiment, there were no differences in mouse body weight or fecal bacterial load based on diet or colonization status (**Extended Data Fig. 1b, c**). Mice monocolonized with *B. thetaiotaomicron* exhibited higher average daily intake of their non-fermentable ID compared to fermentable LD (**Fig. 1f, g**), which was mainly driven by the order of diet exposure: mice exposed to their non-fermentable fructan first (*Bt* NF➔F) ate less of the fermentable fructan compared to the non-fermentable fructan (**Fig. 1h, i**). While no significant differences in diet intake were seen when considering all mice colonized with *B. ovatus* (**Fig. 1f, g**), the same pattern of increased preference for the non-fermentable diet was apparent when groups were examined by diet order: only when exposed to the non-fermentable diet first (*Bo* NF➔F) did mice consume more of their non-fermentable LD than fermentable ID (**Fig. 1h, i**). To further gain insight into whether this difference in diet intake reflects an established dietary preference, mice were subjected at the end of the experimental paradigm to both diets at the same time within the home cage, and overnight intake of each diet was quantified (**Fig. 1e**).

Consistent with results from average daily intake, mice colonized with *B. thetaiotaomicron* ate more of their non-fermentable ID than fermentable LD in the overnight assay (**Fig. 1j, j**), which was mainly driven by the cohort exposed to the non-fermentable fructan first (*Bt* NF➔F) (**Fig. 1l, m**). A similar pattern of preference for the non-fermentable diet, in a diet order-dependent manner, was seen in mice colonized with *B. ovatus*, but it did not reach statistical significance for the overnight assay (**Fig. 1j-m**). Principal component analysis of aggregate data for average daily dietary intake and overnight dietary intake revealed apparent separation of individuals by diet exposure order along PC1, with clearer separation for the *B. thetaiotaomicron* group (**Extended Data Fig. 1d**). The stronger phenotypes with *B. thetaiotaomicron* may be due to its clearer growth bias in LD than ID, whereas *B. ovatus* exhibited substantial baseline growth in LD that was only modestly surpassed in ID (**Fig. 1c**). Overall, the results strongly suggest that gut microbes with differential fructan utilization direct host diet preference toward non-fermentable fructans.

### Bacterial fructan utilization drives host diet preference for non-fermentable diet

Fructan PUL components *susC* and *susD* determine binding specificity and utilization of inulin^22,23^ and levan^20,21^ (**Extended Data Fig. 2a**). To determine whether gut bacterial fructan utilization is necessary or sufficient to alter host diet preference, we used allelic exchange^24^ to manipulate the fructan utilization ability of *B. thetaiotaomicron* and *B. ovatus*. We deleted the *susC*- and *susD*-like genes in the fructan PUL of *B. thetaiotaomicron* (BT1762-1763) and *B. ovatus* (BACOVA_04504-04505) to generate *Bt*^Δ*susCD*^ and *Bo*^Δ*susCD*^, which were no longer able to grow in minimal media containing inulin or levan (**Fig. 2a**, **3a**). To generate an isogenic strain of *B. thetaiotaomicron* that can degrade inulin but not levan, we replaced BT1762-1763 with BACOVA_04504-04505, and deleted the levan-specific glycoside hydrolase BT1760 (**Extended Data Fig. 2c**). The resulting strain *Bt^Bo-susCD^*exhibited the fructan specificity of *B. ovatus*, growing in minimal media containing inulin but not levan (**Extended Data Fig. 2a**). We further confirmed that the genetic modifications resulted in functional differences in production of SCFAs^12^. *B. thetaiotaomicron* produced high levels of acetate when grown in its fermentable LD, which was substantially diminished in *Bt*^Δ*susCD*^ , and reversed in *Bt^Bo-susCD^* which produced high levels of acetate when grown in ID (**Extended Data Fig. 2d**). This aligned with results from *B. ovatus*, where the wild-type strain produced high acetate levels when grown in its fermentable ID, but *Bo*^Δ*susCD*^ failed to produce acetate in either ID or LD. These patterns were not seen for propionate (**Extended Data Fig. 2e**), suggesting that acetate is the dominant SCFA produced by fructan utilization for these particular bacterial species.

**Figure 2:**
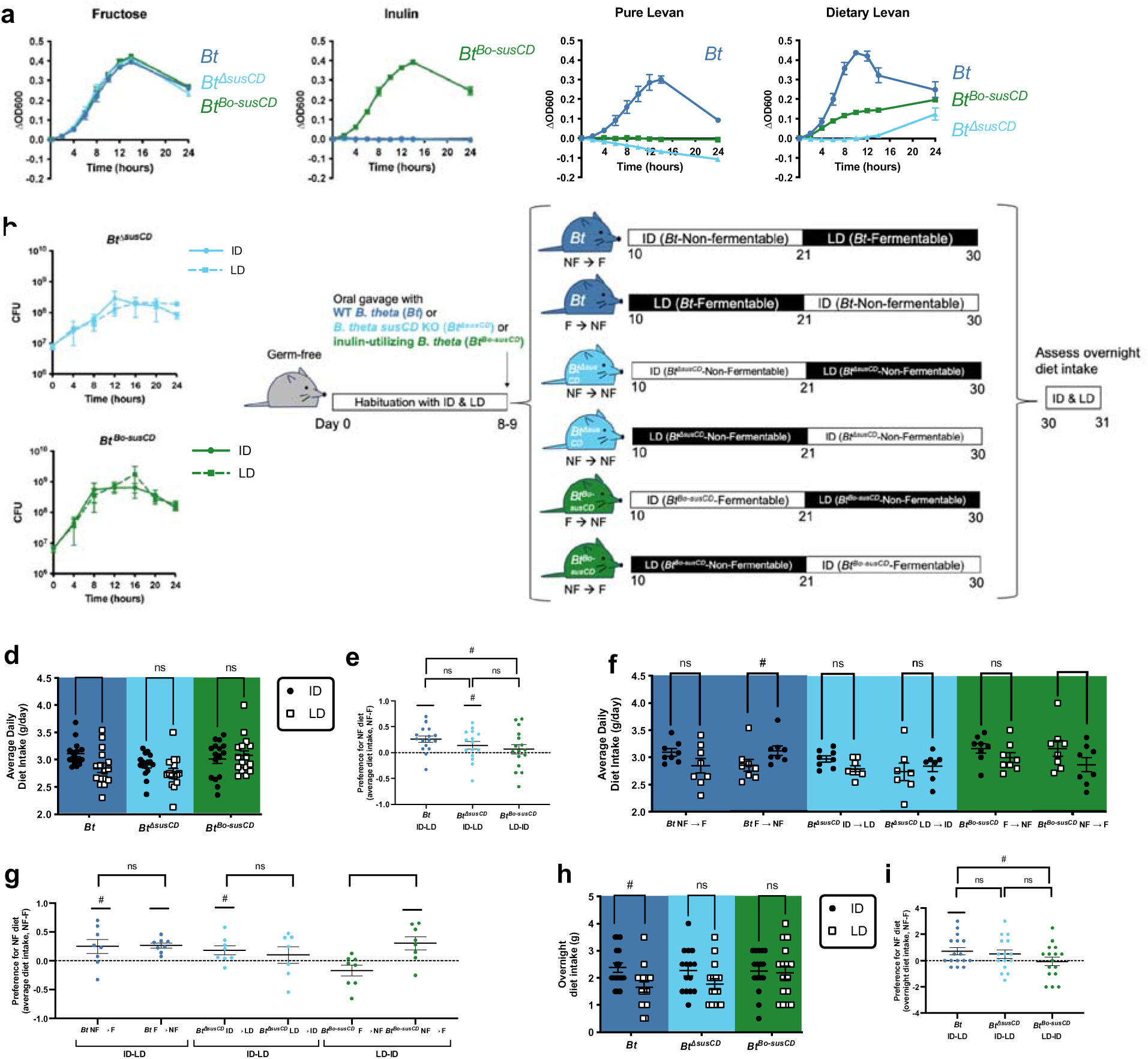
Bacterial fructan utilization drives host preference for diets containing the non-fermentable fructan. **a**) Growth curves of wild-type *B. thetaiotaomicron (Bt)*, mutant *B. thetaiotaomicron* lacking *susCD* (*Bt*^Δ*susCD*^), and inulin-utilizing *B. thetaiotaomicron* (*Bt^Bo-susCD^*) in minimal media containing 0.5% fructose, inulin, levan (pure), or levan (purified for dietary formulation). Each dot represents 3 biological replicates. **b**) Growth curves of *Bt*, *Bt*^Δ*susCD*^, and *Bt^Bo-susCD^* in minimal media containing 5% inulin diet (ID) or levan diet (LD). Each dot represents 3 biological replicates. **c**) Experimental schematic for sequential feeding post-colonization. **d**) Average daily diet intake post-colonization for mice colonized with *Bt*, *Bt*^Δ*susCD*^, and *Bt^Bo-susCD^*. **e**) Preference score for the NF fructan in average daily diet intake post-colonization for mice colonized with *Bt* (ID-LD), *Bt*^Δ*susCD*^ (ID-LD), and *Bt^Bo-susCD^*(LD-ID). **f**) Average daily diet intake post-colonization for mice colonized with *Bt*, *Bt*^Δ*susCD*^, and *Bt^Bo-susCD^*by diet order. **g**) Preference score for the NF fructan in average daily diet intake post-colonization for mice colonized with *Bt* (ID-LD), *Bt*^Δ*susCD*^ (ID-LD), and *Bt^Bo-susCD^* (LD-ID) by diet order. **h**) Overnight diet intake from days 30-31 for mice colonized with *Bt*, *Bt*^Δ*susCD*^, and *Bt^Bo-susCD^* . **i)** Preference score for the NF fructan in overnight diet intake for mice colonized with *Bt* (ID-LD), *Bt*^Δ*susCD*^ (ID-LD), and *Bt^Bo-susCD^* (LD-ID). Data from d-i are combined from two independent experiments. For d, e, h, and i, n = 16 for *Bt*, 15 for *Bt*^Δ*susCD*^, and 16 for *Bt^Bo-susCD^*. For f and g, n = 8 for *Bt* NF ➔ F, 8 for *Bt* F ➔ NF, 8 for *Bt*^Δ*susCD*^ ID ➔ LD, 7 for *Bt*^Δ*susCD*^ LD ➔ ID, 8 for *Bt^Bo-susCD^* F ➔ NF, and 8 for *Bt^Bo-susCD^* NF ➔ F. For d, f, and h, 2-way ANOVA with matched column measures (each mouse), comparing means across rows, and Sidak’s corrections were performed. For e, g, and i, 90 and 95% confidence intervals were computed, and significance was determined if the interval did not contain zero. Unpaired parametric t-tests were also performed between groups. # = p-value < 0.10; * = p-value < 0.05; ** = p-value < 0.01

**Figure 3:**
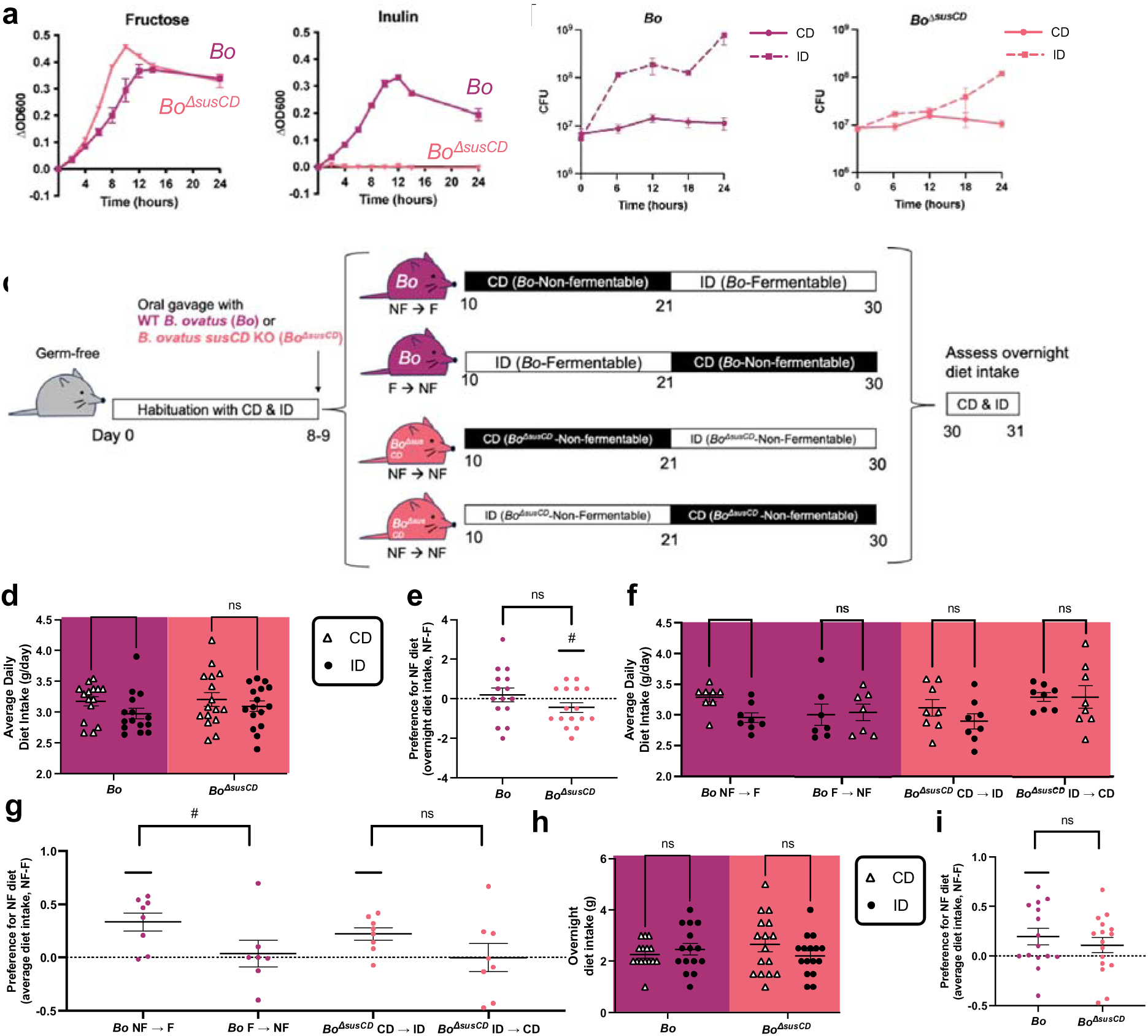
Colonization with fructan-utilizing gut bacteria increases host preference for fructan-free diet. **a**) Growth curves of wild-type *B. ovatus (Bo)* and *B. ovatus susCD* KO (*Bo*^Δ*susCD*^) in minimal media containing 0.5% fructose or inulin. Each dot represents 3 biological replicates. **b**) Growth curves of *Bo* and *Bo*^Δ*susCD*^ in minimal media containing 5% cellulose diet (CD) or inulin diet (ID). Each dot represents 3 biological replicates**. c**) Experimental schematic for sequential feeding post-colonization**. d)** Average daily diet intake post-colonization for mice colonized with *Bo* and *Bo*^Δ*susCD*^. **e**) Preference score for the NF fructan in average daily diet intake post-colonization for mice colonized with *Bo* (CD-ID) and *Bo*^Δ*susCD*^ (CD-ID). **f**) Average daily diet intake post-colonization for mice colonized with *Bo* and *Bo*^Δ*susCD*^ by diet order. **g**) Preference score for the NF fructan in average daily diet intake post-colonization for mice colonized with *Bo* (CD-ID) and *Bo*^Δ*susCD*^ (CD-ID) by diet order. **h**) Overnight diet intake from days 30-31 for mice colonized with *Bo* and *Bo*^Δ*susCD*^. **i**) Preference score for the NF fructan in overnight diet intake for mice colonized with *Bo* (CD-ID) and *Bo*^Δ*susCD*^ (CD-ID). Data from d-i are combined from two independent experiments. For d, e, h, and i, n = 15 for *Bo*, 16 for *Bo*^Δ*susCD*^ . For f and g, n = 8 for *Bo* NF ➔ F, 7 for *Bo* F ➔ NF, 8 for *Bo*^Δ*susCD*^ CD ➔ ID, and 8 for *Bo*^Δ*susCD*^ ID ➔ CD. For d, f, and h, 2-way ANOVA with matched column measures (each mouse), comparing means across rows, and Sidak’s corrections were performed. For e, g, and i, 90 and 95% confidence intervals were computed, and significance was determined if the interval did not contain zero. Unpaired parametric t-tests were also performed between groups. # = p-value < 0.10; * = p-value < 0.05

To determine how bacterial fructan utilization impacts host feeding behavior, we monocolonized mice with wildtype *B. thetaiotaomicron*, deletion mutant *Bt*^Δ*susCD*,^ or gene replacement mutant *Bt^Bo-susCD^*, fed them ID or LD in sequence for 10 days each, and measured their food consumption (**Fig. 2c**). We confirmed that *Bt*^Δ*susCD*^ exhibits poor growth in minimal media containing ID or LD (**Fig. 2a, b**), as compared to wildtype *B. thetaiotaomicron* (**Fig. 1b, c**). *Bt^Bo-susCD^*exhibited strong growth in minimal media containing inulin and ID, but like *B. ovatus* (**Fig. 1b, c**), it had no growth with pure levan and intermediate growth with dietary levan and LD (**Fig. 2a, b**, **Extended Data Fig. 3a**). Mice showed no differences in body weight or bacterial load based on diet or colonization status (**Extended Data Fig. 3b, c**), indicating that fructan utilization is not required for successful gut colonization. As previously observed (**Fig. 1e-h**), mice colonized with wild-type *B. thetaiotaomicron* exhibited higher average daily intake of their non-fermentable ID compared to fermentable LD (**Fig. 2d-g**). They also exhibited increased preference for their non-fermentable ID when given the choice between ID and LD in an overnight feeding assay (**Fig. 2h, i**). These phenotypes were abrogated in mice colonized with *Bt*^Δ*susCD*^, which exhibited no statistical differences in average daily food intake or preference for ID vs. LD (**Fig. 2d-g**). This suggests that bacterial fructan utilization is necessary to mediate host preference for diets containing the non-fermentable fructan. Average daily intake was further altered by colonization with *Bt^Bo-susCD^* only in a manner dependent on the order of diet exposure: *Bt^Bo-susCD^* NF➔F mice ate less of the fermentable ID compared to the non-fermentable LD (**Fig. 2f, g**); this modest change is comparable to the effect seen with *B. ovatus* (**Fig. 1g, h**) and may again be due to the notable baseline growth of *Bt^Bo-susCD^* in dietary levan and LD, relative to inulin and ID (**Fig. 2a, b**). In an overnight assay, mice colonized with *Bt* ate more non-fermentable ID than fermentable LD, while mice colonized with *Bt*^Δ*susCD*^ or *Bt^Bo-susCD^* showed no difference (**Figure 2h, i**, **Extended Data Fig. 3d, e**). Together, these results indicate that bacterial fructan utilization is necessary and at least partially sufficient to alter host food consumption and food choice behavior toward diets containing the non-fermentable fructan.

To confirm that bacterial fructan utilization is necessary to alter host food preference, we next tested the effect of *Bo*^Δ*susCD*^, which cannot grow on inulin (**Fig. 3a**), compared to wild-type *B. ovatus*. Due to the observed baseline growth due to presumed impurities in the dietary levan and non-fermentable LD (**Fig. 1a, b**), we switched to a 10% cellulose diet (CD) which contains cellulose as its only carbohydrate source to serve as the *B. ovatus*-non-fermentable diet. As expected, *B. ovatus* exhibits growth in minimal media containing ID, but not CD (**Fig. 3b**, **Extended Data Fig. 4a**), which reflects a clear growth differential that was not as apparent when using LD as the non-fermentable diet (**Fig. 1c**). Furthermore, *B. ovatus* produced more acetate and propionate when growing on fermentable ID compared to non-fermentable CD (**Extended Data Fig. 2f, g**) to a greater degree than when growing on fermentable ID compared to non-fermentable LD (**Extended Data Fig. 2d, e**). Growth and SCFA production in ID were substantially decreased in the deletion strain *Bo*^Δ*susCD*^ (**Fig. 3b**, **Extended Data Fig. 4a**, **Extended Data Fig. 2h, i**), so we hypothesized that colonization with *Bo*^Δ*susCD*^ would diminish host preference for the non-fermentable diet seen with *B. ovatus* colonization. To test this, we monocolonized mice with *B. ovatus* or *Bo*^Δ*susCD*^, fed them CD or ID in sequence for 10 days each, and measured their food consumption (**Fig. 3c**). There were no differences in mouse body weight based on diet or colonization status (**Extended Data Fig. 4b**). There was a slight decrease in bacterial load in mice when switching from fermentable ID to non-fermentable CD, suggesting that *B. ovatus* has a growth disadvantage on cellulose but still colonizes the intestine (**Extended Data Fig. 4c**). Consistent with our prior observations with ID and LD (**Fig. 1g, h**), mice colonized with *B. ovatus* exhibited higher average daily intake of the non-fermentable CD compared to the fermentable ID (**Fig 3d, e**), which was seen particularly in mice exposed in the NF➔F diet order (**Fig 3f, g**). This effect was abrogated or diminished in mice colonized with deletion strain *Bo*^Δ*susCD*^ (**Fig. 3d-g**), again suggesting that bacterial fructan utilization mediates differential host diet intake. However, there were no differences between *B. ovatus*-colonized groups in overnight diet intake (**Fig. 3h, i**, **Extended Data Fig. 4d, e**), which differs from mice colonized with *B. thetaiotaomicron* (**Fig. 2h, i**). All together, these results indicate that bacterial fructan utilization is necessary to modulate host preference for diets containing the non-fermentable fructan.

**Figure 4:**
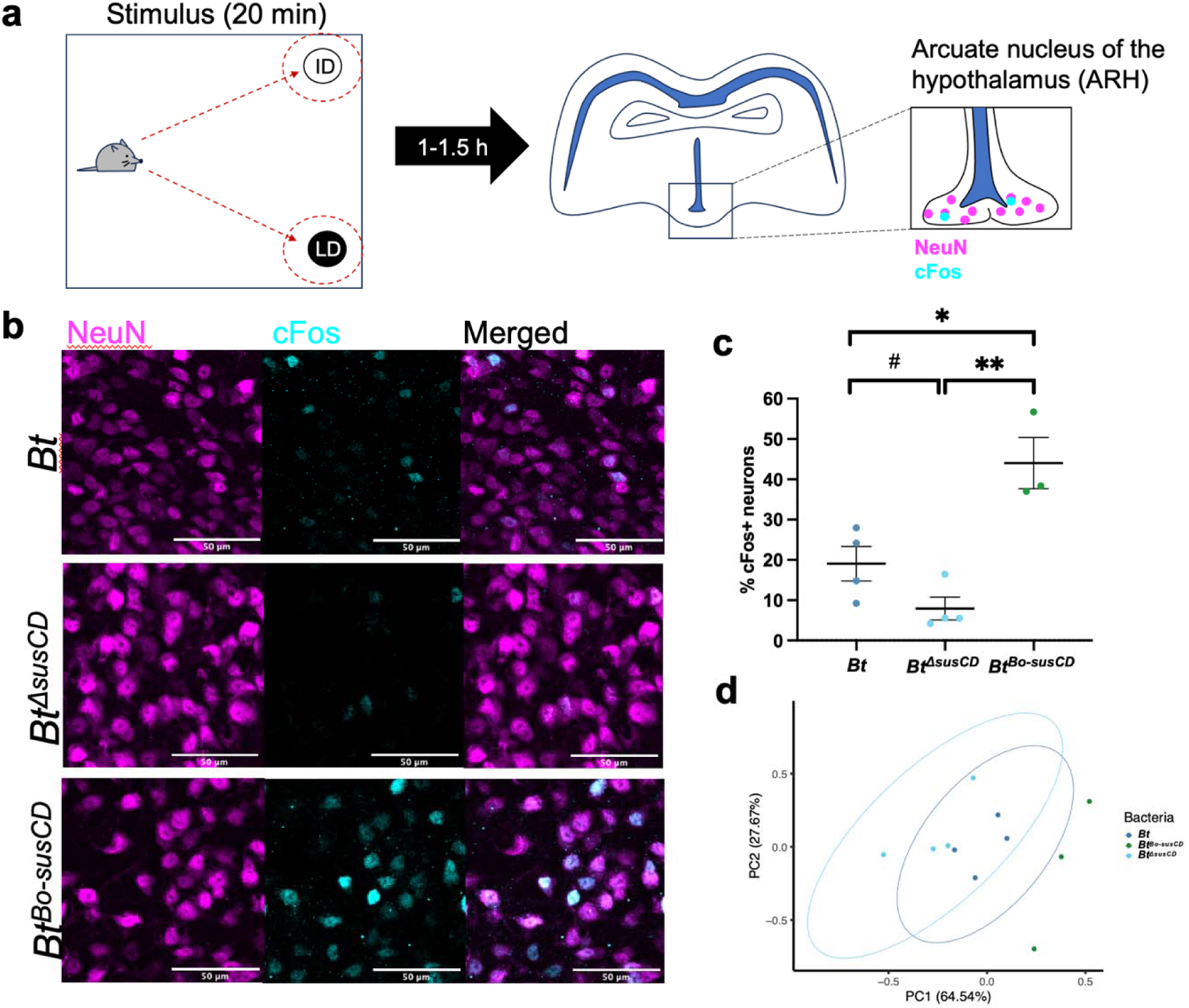
Bacterial fructan utilization by *B. thetaiotaomicron* promotes diet preference-induced neuronal activation in the arcuate nucleus of the hypothalamus. **a)** Schematic of experimental set up: 1-1.5 hours after the fasting-induced, mice were sacrificed, and their brains collected. Brains were sectioned coronally for the arcuate nucleus of the hypothalamus (ARH), and ARH sections were stained for neurons (NeuN, magenta) and early activation marker, cFos (cyan). **b**) Representative figures of ARH in mice colonized with *Bt*, *Bt*^Δ*susCD*^, and *Bt^Bo-susCD^*. **c)** Quantification of cFos-positive neurons in ARH of mice colonized with *Bt*, *Bt*^Δ*susCD*^, and *Bt^Bo-susCD^* (n = 3-4 mice per group). Each dot represents the average of three technical replicates. **d**) PCA plots for average daily diet intake, overnight diet intake, and cFos % in mice colonized with *Bt*, *Bt*^Δ*susCD*^ , and *Bt^Bo-susCD^* (n = 3-4 mice per group). For c, unpaired parametric t-tests were performed. # = p-value < 0.10; * = p-value < 0.05; ** = p-value < 0.01

### Bacterial fructan utilization induces neuronal activation in feeding-related brain region after food choice

Neurons in the arcuate nucleus of the hypothalamus (ARH) are important regulators of homeostatic feeding behavior^25,26^ that respond rapidly to intragastric nutrient infusion^27^ and mediate food associations with sensory or environmental cues^26,28,29^. Neurons in ARH, but not other hypothalamic regions, are activated in response to intraperitoneal injection of acetate^30^. We thus hypothesized that host preference for diets containing the non-fermentable fructan reflect association of fermentable diet with metabolites generated by bacterial fructan utilization, which may be encoded by neuronal activity in the ARH. To test whether bacterial fructan utilization impacts neuronal activity in the ARH, we examined cFos as an early neuronal activation marker^31,32^ in mice pre-exposed to the sequential feeding paradigm for associative learning and then subjected to both diets to trigger acute food preference as a stimulus on day 32 (**Fig. 4a**). In order to limit potential confounding factors, mice were fasted overnight and then habituated to a sterile open field arena before exposure to both diets. Mice colonized with *B. thetaiotaomicron* or *Bt^Bo-susCD^*, which exhibited host preference for diets containing the non-fermentable fructan (**Fig. 1, 2**), exhibited a higher percentage of cFos-positive neurons in the ARH than mice colonized with *Bt*^Δ*susCD*^ (**Fig. 4b, c**), which did not exhibit host dietary preference (**Fig. 2**). This suggests that bacterial fructan utilization is necessary to promote diet preference-induced neuronal activation in the ARH, which aligns with host behavioral preference for consuming diets containing the non-fermentable fructan (**Fig. 1, 2**). Principal component analysis of data for diet preference-induced neuronal activation in the ARH, average daily diet intake, and overnight diet intake discriminates mice colonized with *B. thetaiotaomicron* from those colonized with *Bt*^Δ*susCD*^ or *Bt^Bo-susCD^*along PC1 (**Fig. 4d**), suggesting that bacterial fructan utilization co-regulates dietary preference and ARH neuronal activation. Overall, results from this study reveal a role for bacterial fructan utilization in conditioning host preference for fructan-containing diets and associated neuronal response in the ARH.

## Discussion

This study demonstrates that selective nutrient utilization by gut bacteria can modulate host diet preferences. We focused on bacterial metabolism of two fructans —inulin and levan— that differ only in their glycosidic linkages and thus are expected to exhibit similar perceptual salience to the host. This contrasts prior studies that have examined microbial influences on host preference for palatable versus non-palatable diets that differ substantially in macronutrient content^8^. Results from our experiments reveal that even with equivalent formulation of different fructan-containing diets, fructan-utilizing gut bacteria direct the host to increase relative consumption of diets containing the non-fermentable fructan. This effect of bacterial fructan utilization on host dietary fructan preference was reproduced using two prevalent bacteria of the human gut microbiome – *B. thetaiotaomicron* and *B. ovatus*. Interestingly, the phenotypes observed with *B. ovatus* were consistently weaker than those observed with *B. thetaiotaomicron*, as was the efficiency of fructan utilization by *B. ovatus* compared to *B. thetaiotaomicron.* The results suggest that the strength of host diet association may depend on the efficiency of bacterial fructan degradation and amount of its fermentation products, such as the SCFA acetate, to the host. *B. thetaiotaomicron* grows robustly on its fermentable LD, reaching over 10^9^ colony forming units by hour 12 (**Fig. 1c**, **Extended Data Fig. 3a**), while *B. ovatus* does not reach 10^9^ colony forming units by hour 24 when grown on its fermentable ID (**Fig. 1c**, **Extended Data Fig. 4a**), suggesting that *B. thetaiotaomicron* is more efficient than *B. ovatus* at utilizing fructans. Furthermore, *B. thetaiotaomicron* produces significantly more acetate on its fermentable LD compared to non-fermentable ID than *B. ovatus* does on its fermentable ID compared to non-fermentable LD (**Extended Data Fig. 2h**). Beyond SCFAs, other metabolites, such lactate and succinate, are also produced by bacterial fermentation of fructans^12^. Further research is warranted to identify the specific signaling molecules that mediate the ability of bacterial fructan utilization to condition host dietary fructan preference.

Experiments in this study employ a reductionist approach with monocolonized mice and engineered bacteria to demonstrate host associative learning of microbiome- and diet-dependent signals in the absence of overt differences in diet palatability. In our sequential feeding paradigm, 10-day exposure to each diet was sufficient for such associative learning to occur. This aligns with a previous report that mice colonized with microbiota from herbivorous wild rodents showed greater preference for diet with a higher protein to carbohydrate ratio, but only after one week^7^. We observed that associative learning appears stronger depending on the order of diets that mice were exposed to: mice exposed to the non-fermentable diet first show a stronger preference for non-fermentable diet than those exposed to fermentable diet first. The host preference for non-fermentable diet seems initially counterintuitive because products of bacterial fermentation such as SCFAs can serve as additional energy sources for host intestinal epithelial cells^12^, which means more energy can be extracted from the diet.

This extra energy along with the known role of SCFAs in reducing host appetite^12–15^ may lead the host to eat less fermentable diet and associate the fermentable diet with increased satiety, only if they are exposed to non-fermentable diet first to have a comparative baseline. One potential confound may be intestinal gas and bloating resulting from bacterial fructan fermentation, but a regular laboratory mouse diet includes ∼15% soluble plant fibers^33^, so we do not expect our 10% fructan diet to cause more fiber-associated discomfort than a standard diet, and we did not observe weight loss or other gross appearance or behavioral differences in mice consuming a fermentable diet. From an evolutionary perspective, host preference for non-fermentable diet over fermentable diet may keep bacterial loads in check through a negative feedback loop of bacterial satiety signals such as SCFAs by preventing a bloom of bacteria when they can utilize additional nutrient sources. This is supported by our observation that bacterial loads are not significantly different on fermentable or non-fermentable diets (**Extended Data Fig. 1c, 3c**). Our finding aligns with a previous report in which intestinal *E. coli* shows exponential growth *in vivo* in response to nutrient infusions and produces satiety proteins such as ClpB that activate anorexigenic POMC neurons in the ARH^34^.

Consistent with the observation that the bacterial protein ClpB can act on ARH neurons^34^, we found that bacterial fructan utilization promotes diet preference-induced neuronal activation in the ARH. We hypothesize that metabolites from bacterial fructan utilization may act on the ARH to promote satiety, which over the 10-day exposure period, may lead mice to associate the fermentable diet with satiety in a diet order-dependent manner; thus, mice would consume less fermentable diet on average and consume more non-fermentable diet when presented both. AgRP neurons in the hypothalamus promote hunger and lead to negative-valence learning in which mice avoid AgRP activation ^29^. If bacterial metabolites activate AgRP neurons in the ARH, such negative-valence learning may explain mice’s preference for non-fermentable over fermentable diet. However, future studies are needed to parse whether the neuronal activation causes or is the result of diet preference behavior. Furthermore, the mechanisms by which this occurs warrant further investigation. We propose metabolites from fructan utilization may act on intestinally innervating afferent vagal neurons or travel through the circulation to have direct effects on the brain. For example, one study reports the ability of the SCFA acetate to cross the blood-brain-barrier to act on hypothalamic neurons to suppress appetite^35^, but other microbial mechanisms remain possible. The vagal afferent neurons play a key role in nutrient sensing and preference^36^, such as modulating nutrient preference for sugar over artificial sweetener^32,37^. It also expresses many microbial metabolite receptors^38,39^ and connects to feeding-related brain regions such as the ARH^40^ and striatum^41,42^ through multi-synaptic pathways. Through selective ablation of gut-innervating vagal neurons, a role for vagal nerve signaling has also been shown for bacterial modulation of high fat diet preference^9^. However, most literature studying bacterial modulation of host diet preference has focused on preference for palatable diets and reward-related brain regions such as the nucleus accumbens^8–10^. The role of bacterial modulation of homeostatic feeding circuits in the ARH through the vagus nerve remains unclear. Traditionally associated with satiety, AgRP neurons are increasingly appreciated for their involvement in flavor nutrient learning^28^, highlighting the importance of understanding how bacteria can modulate ARH activity. Understanding the mechanisms by which gut microbes influence host food choice could potentially inform new approaches for promoting healthier eating habits and ameliorating metabolic and eating disorders.

## Acknowledgements

We thank members of the Hsiao laboratory for their guidance and review of the manuscript; members of the UCLA Goodman-Luskin Microbiome Center Gnotobiotics Core Facility for protocol development and technical support; Dr. Jieping Yang, Dr. Ru- Po Lee, and Dr. Qing-Yi Lu of the UCLA Analytical Phytochemical Core for measuring SCFAs; Dr. Karen Reue and Dr. Laurent Vergnes for conducting pilot investigations of mitochondrial metabolism; Dr. Andrew Goodman for providing the pSIE1 plasmid; and Dr. Joan Combie (Montana Biopolymers Corp.) for producing levan and advising on its purification. This work was supported by funds from a UCLA-Caltech Medical Scientist Training Program T32 (NIGMS 3T32GM008042-35S1) and UCLA Whitcome Fellowship to K.B.Y. and W.M. Keck foundation grant to E.Y.H. E.Y.H. is a New York Stem Cell Foundation – Robertson Investigator. This research was supported in part by the New York Stem Cell Foundation.

## Author Contributions

K.B.Y., C.S., A.C., J.P., A.N., E.O., S.A.K., G.R.L., and A.L. performed the experiments and analyzed the data. E.O. and J.B.L. provided key technical guidance and resources. K.B.Y. and E.Y.H. designed the study and wrote the manuscript. All authors discussed the results and commented on the manuscript.

## Extended Data Figures and Figure Legends

**Extended Data Figure 1:**
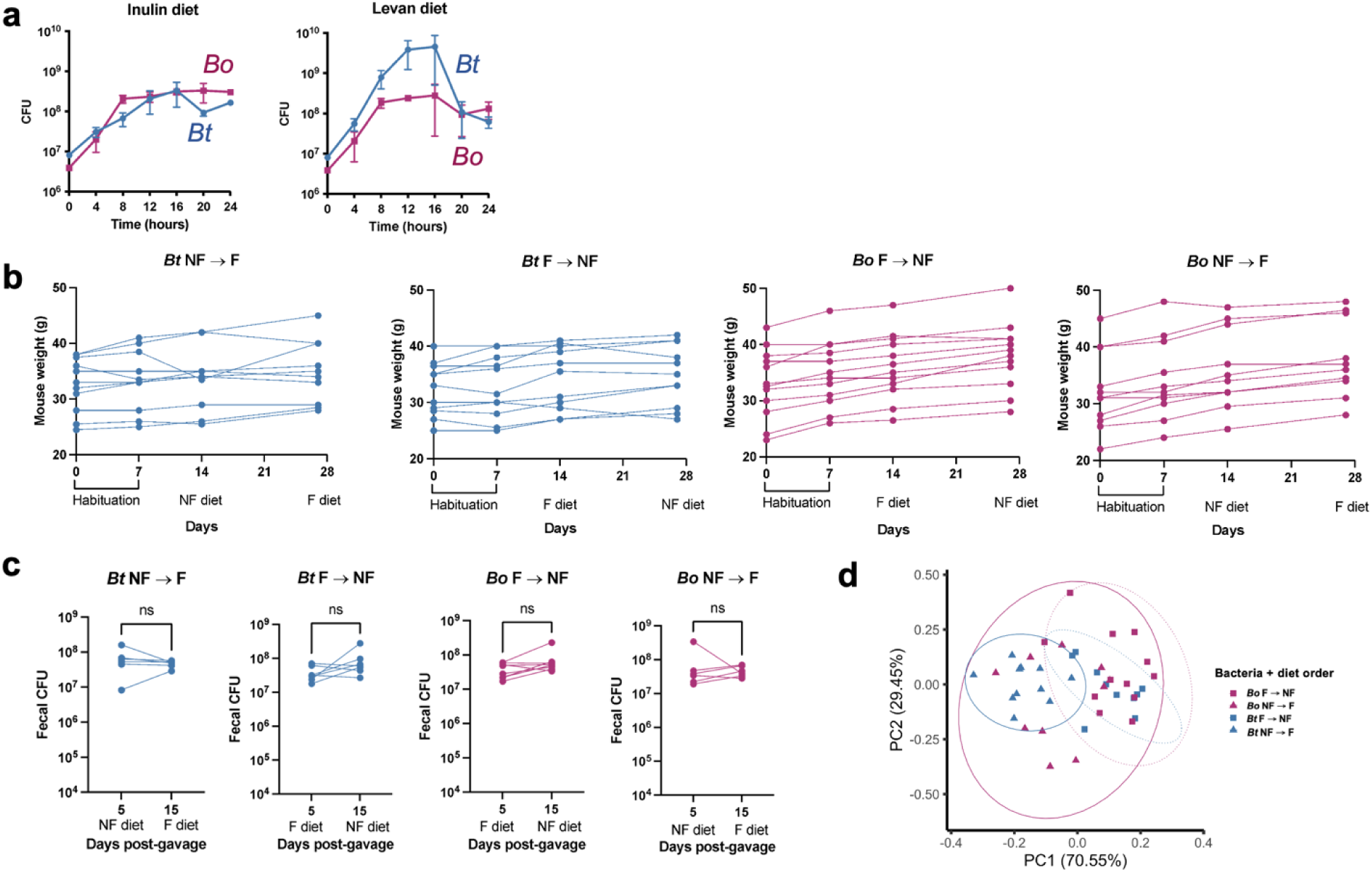
Body weight and bacterial load in *Bacteroides* mono-colonized mice fed diets containing non-fermentable or fermentable fructan. **a)** Growth curves of *Bt* and *Bo* in minimal media containing 5% inulin diet (ID) or levan diet (LD). **b)** Mouse weights during the sequential feeding experiment, separated by bacteria and diet order. The diets that mice were consuming are listed underneath the x-axis. **c**) Mouse fecal bacterial load during the sequential feeding experiment, separated by bacteria and diet order. The diets that mice were consuming are listed underneath the x-axis. **d**) PCA plot for average daily diet intake and overnight diet intake. Solid ellipses surround NF➔F, while dotted ellipses surround F➔NF. Data for b-d are combined from three independent experiments. For b and d, n = 11 for *Bt* NF ➔ F, 11 for *Bt* F ➔ NF, 12 for *Bo* F ➔ NF, 11 for *Bo* NF ➔ F. For c, n = 6 for *Bt* NF F, 7 for *Bt* F ➔ NF, 8 for *Bo* F ➔ NF, 6 for *Bo* NF ➔ F. For c, paired parametric t-tests were performed.

**Extended Data Figure 2:**
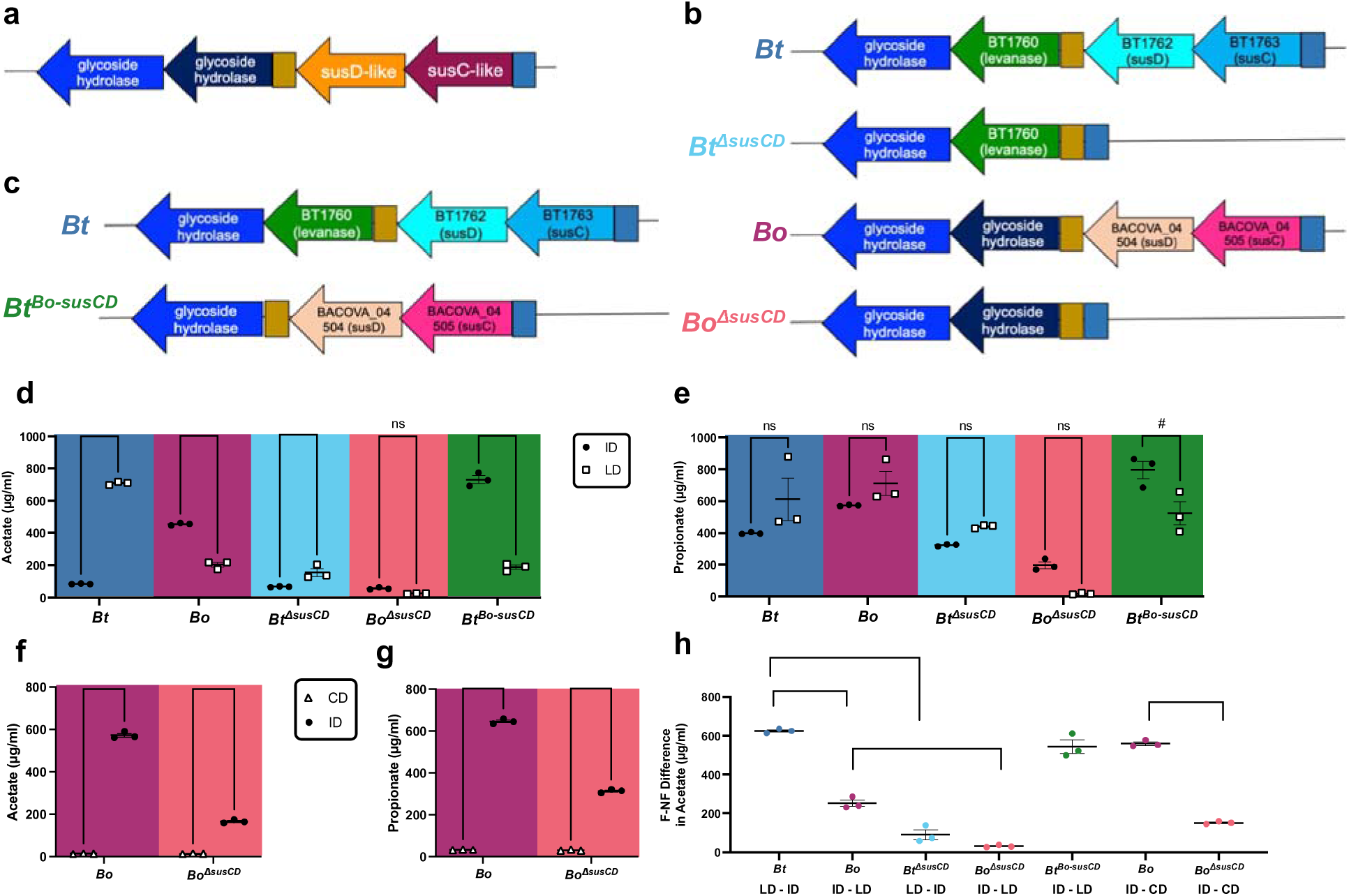
Validation of engineered *Bacteroides* in growth assays and production of short chain fatty acids. **a**) Schematic of fructan *sus* operon in both *Bt* and *Bo*. One of the glycoside hydrolases in *Bt* is specific for linkages in levan, called levanase (BT1760). *susC-* and *susD-like* genes from this operon are henceforth referred to as *susC* and *susD* or *susCD* genes. **b**) Schematic of fructan *sus* operons for Δ*susCD* bacterial strains. **c**) Schematic of fructan *sus* operon for inulin-utilizing *B. thetaiotomicron (Bt^Bo-susCD^*). **d**) Acetate produced in supernatants of bacteria grown for 24 hours in minimal media containing 5% inulin diet (ID) or levan diet (LD). (n = 3 each) Propionate produced in supernatants of bacteria grown for 24 hours in minimal media containing 5% ID or LD. (n = 3 each) **f**) Acetate produced in supernatants of bacteria grown for 24 hours in minimal media containing 5% cellulose diet (CD) or ID. (n = 3 each) **g**) Propionate produced in supernatants of bacteria grown for 24 hours in minimal media containing 5% CD or ID. (n = 3 each) **h**) Difference (F-NF) in acetate produced in bacterial supernatants grown on fermentable (F) and non-fermentable (NF) diet. Diets are listed underneath the x-axis. (n = 3 each). For d-g, 2-way ANOVA with matched column measures (each bacterial colony), comparing means across rows, and Sidak’s corrections were performed. For h, unpaired parametric t-tests were performed. # = p-value < 0.10, * = p-value < 0.05; ** = p-value < 0.01; *** = p-value < 0.001; **** = p-value < 0.0001

**Extended Data Figure 3:**
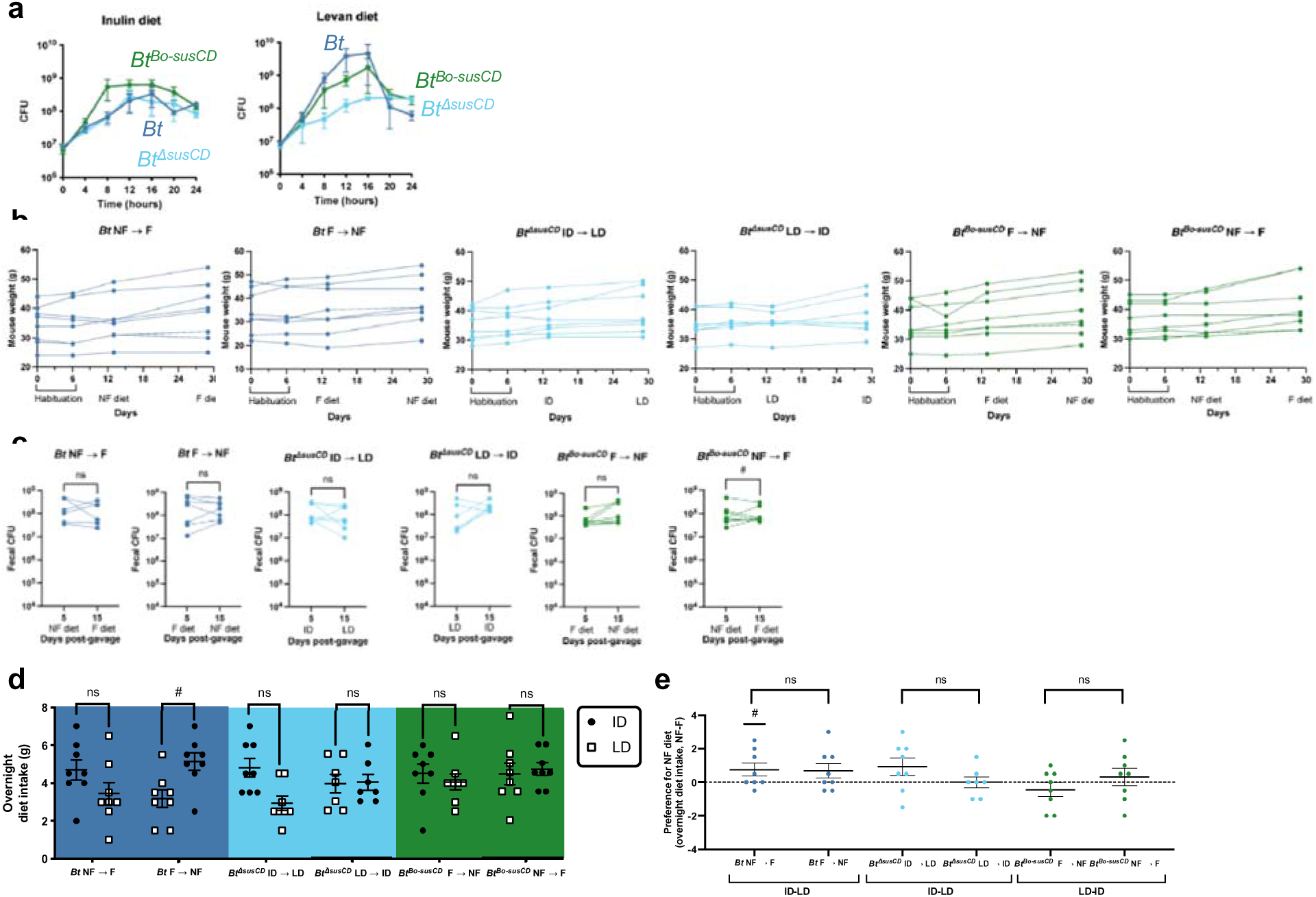
Body weight, bacterial load, and overnight diet intake in mice mono-colonized with wildtype or engineered *B. thetaiotaomicron*. **a)** Growth curves of *Bt*, *Bt*^Δ*susCD*^, and *Bt^Bo-susCD^* in minimal media containing 5% inulin diet (ID) or levan diet (LD). Each dot represents 3 biological replicates. **b**) Mouse weights during the sequential feeding experiment, separated by bacteria and diet order. The diets that mice were consuming are listed underneath the x-axis. **c**) Mouse fecal bacterial load during the sequential feeding experiment, separated by bacteria and diet order. The diets that mice were consuming are listed underneath the x-axis. **d**) Overnight diet intake from days 30-31 for mice colonized with *Bt*, *Bt*^Δ*susCD*^, and *Bt^Bo-susCD^* by diet order. **e**) Preference score for the NF fructan in overnight diet intake for mice colonized with *Bt* (ID-LD), *Bt*^Δ*susCD*^ (ID-LD), and *Bt^Bo-susCD^* (LD-ID) by diet order. Data from b-e are combined from two independent experiments. For b, d, and e, n = 8 for *Bt* NF ➔ F, 8 for *Bt* F ➔ NF, 8 for *Bt*^Δ*susCD*^ ID ➔ LD, 7 for *Bt*^Δ*susCD*^ LD ➔ ID, 8 for *Bt^Bo-susCD^* F ➔ NF, and 8 for *Bt^Bo-susCD^* NF ➔ F. For c, n = 6 for *Bt* NF ➔ F, 7 for *Bt* F ➔ NF, 7 for *Bt*^Δ*susCD*^ ID ➔ LD, 6 for *Bt*^Δ*susCD*^ LD ➔ ID, 8 for *Bt^Bo-susCD^* F ➔ NF, and 8 for *Bt^Bo-susCD^* NF ➔ F. For c, paired parametric tests were performed. For d, 2-way ANOVA with matched column measures (each mouse), comparing means across rows, and Sidak’s corrections were performed. For e, 90 and 95% confidence intervals were computed, and significance was determined if the interval did not contain zero. Unpaired parametric t-tests were also performed between groups. # = p-value < 0.10; * = p-value < 0.05

**Extended Data Figure 4:**
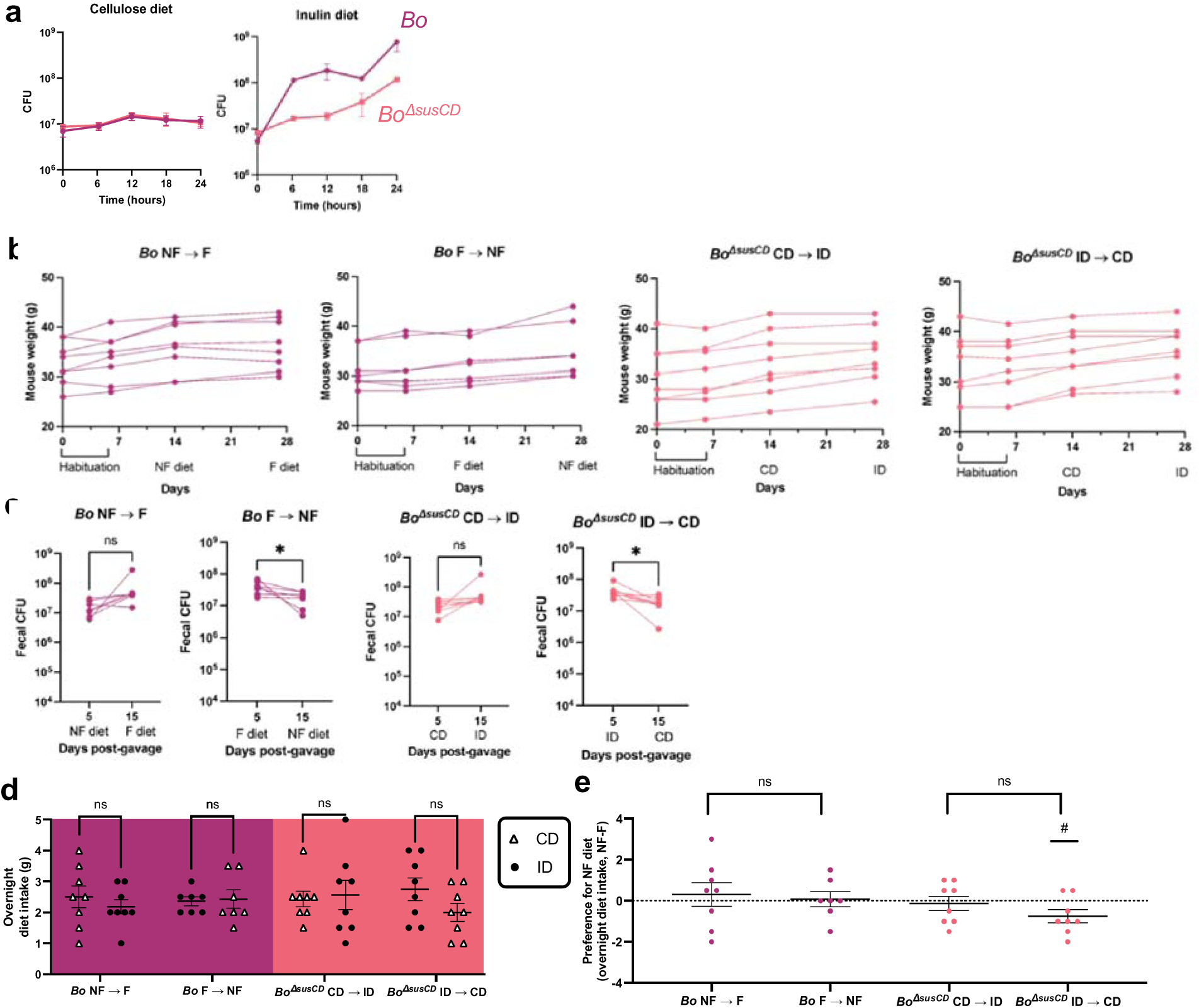
Body weight, bacterial load, and overnight diet intake in mice mono-colonized with wildtype or engineered *B. ovatus*. **a)** Growth curves of *Bo* and *Bo*^Δ*susCD*^ in minimal media containing 5% cellulose diet (CD) or inulin diet (ID). Each dot represents 3 biological replicates. **b**) Mouse weights during the sequential feeding experiment, separated by bacteria and diet order. The diets that mice were consuming are listed underneath the x-axis. **c**) Mouse fecal bacterial load during the sequential feeding experiment, separated by bacteria and diet order. The diets that mice were consuming are listed underneath the x-axis. **d**) Overnight diet intake from days 30-31 for mice colonized with *Bo* and *Bo*^Δ*susCD*^ by diet order. **e**) Preference score for the NF fructan in overnight diet intake for mice colonized with *Bo* (CD-ID) and *Bo*^Δ*susCD*^ (CD-ID) by diet order. Data from b-e are combined from two independent experiments. For b, d, and e n = 8 for *Bo* NF ➔ F, 7 for *Bo* F ➔ NF, 8 for *Bo*^Δ*susCD*^ CD ➔ ID, and 8 for *Bo*^Δ*susCD*^ ID ➔ CD. For c, n = 7 for *Bo* NF ➔ F, 7 for *Bo* F ➔ NF, 8 for *Bo*^Δ*susCD*^ CD ➔ ID, and 8 for *Bo*^Δ*susCD*^ ID ➔ CD. For c, paired parametric t-tests were performed. For d, 2-way ANOVA with matched column measures (each mouse), comparing means across rows, and Sidak’s corrections were performed. For e, 90 and 95% confidence intervals were computed, and significance was determined if the interval did not contain zero. Unpaired parametric t-tests were also performed between groups. # = p-value < 0.10; * = p-value < 0.05

## Methods

### Culturing Bacteria

*B. thetaiotaomicron* (VPI-5482) and *B. ovatus* (ATCC-8483) were cultured in BHIS— Brain Heart Infusion (BD) supplemented with 5 µg/ml hemin and vitamin K1 (Sigma-Aldrich)—in an anaerobic atmosphere of 85% nitrogen, 10% carbon dioxide, and 5% hydrogen. Growth curves were conducted in Salyers minimal media (MM), which contains 100 mM potassium phosphate buffer (pH 7.2), 7.5 mM (NH_4_)_2_SO_4_, 9.4 mM Na_2_CO_3_, 4mM L-cysteine, 1.4 µM FeSO_4_.7(H_2_O), 1 µg/ml vitamin K3, 5 ng/ml vitamin B_12_, 1.9 µM hematin/200 µM L-histidine, 15 mM NaCl, 180 µM CaCl_2_.2H_2_O, 98 µM MgCl_2_.6H_2_O, 50 µM MnCl_2_.4H_2_O, and 42 µM CoCl_2_.6H_2_O. Fructans (fructose, inulin, levan) were sterilized by UV irradiation for 1 hour and added to MM at a final concentration of 0.5%. Fructan diets (CD, ID, LD), which contain 10% fructan, were added to MM at a 5% for a final fructan concentration of 0.5%. For all growth curves, single colonies of bacteria were grown overnight in liquid BHIS, sub-cultured 1:1000 to synchronize their growth, washed with MM, and sub-cultured 1:25 into MM plus fructan or fructan diet for 24 hours. Growth curves with MM plus fructan were obtained by sampling 200 µl from each tube and reading OD600 with a Synergy H1 plate reader at each timepoint. Due to the texture of fructan diets, OD600 could not be read for MM plus fructan diet. Growth curves with MM plus fructan diet were obtained by sampling 10 µl from each tube, performing serial dilutions, and plating 10 µl of each serial dilution on BHIS plates to obtain colony forming units (CFU) at each timepoint.

### Bacterial Genetic Engineering

Gene deletions for *Bt*^Δ*susCD*^ and *Bo*^Δ*susCD*^ mutants were generated using counterselectable allelic exchange with pSIE1 (Addgene plasmid #136355) as described previously^24^. Regions upstream and downstream of the *susC* and *susD* genes in the fructan PUL of *B. thetaiotaomicron* (BT1762-1763) and *B. ovatus* (BACOVA_04504-04505) were PCR amplified (**Table S2**), ligated into the pSIE1 vector via Gibson assembly (New England Biolabs), and conjugated into *Bt* via *E. coli* S17 λ- pir and *Bo* via *E. coli* WM3064. *Bacteroides* colonies that had uptaken the vector were selected with gentamicin (200 µg/ml) and erythromycin (12.5 µg/ml), and counterselection was performed with anhydrotetracycline (100 ng/ml). After two rounds of counterselection, the desired mutants were identified by PCR screening. An isogenic strain of *B. thetaiotaomicron* that can degrade inulin but not levan (*Bt^Bo-susCD^*) was similarly generated by counterselectable allelic exchange to replace BT1762-1763 with BACOVA_04504-04505 (**Table S2**). Desired mutants then underwent counterselectable allelic exchange again to delete the levan-specific glycoside hydrolase BT1760 (**Table S2**). All mutants were confirmed by PCR screening and functional test by growth curves in MM plus fructan.

At 24 hours after inoculation into MM plus fructan diet, 500 µl of bacterial culture was spun down at 16000xg for 2 minutes to obtain cell-free supernatant. Acetate and propionate were measured in cell-free supernatant via gas chromatography performed by the UCLA Analytical Phytochemical Core.

### Sources and Preparation of Fructans

Fructose (F3510), chicory inulin (I2255), and pure levan (L8647) were obtained from Sigma-Aldrich. Dietary levan from Montana Biopolymers Corp. was purified by sequential water and ethanol washes to isolate higher molecular weight fructans and to remove mono- and oligosaccharides. Briefly, 200 grams of crude levan were washed with ddH_2_O for two hours and precipitated using -20° ethanol twice. The resulting levan was then washed with 100% ethanol three times, fully dissolved in ddH_2_O at 75°C, filtered through Whatman paper to remove debris, and dried at 37°C until it became a white powder.

### Mice

All experimental procedures were carried out in accordance with US NIH guidelines for the care and use of laboratory animals and approved by the UCLA Institutional Animal Care and Use Committees. Mice used for data collection were male and female germ-free (GF) wild-type Swiss Webster mice, at least 7-8 weeks of age. GF Swiss Webster mice were purchased from Taconic Farms and bred in flexible film isolators at the UCLA Goodman-Luskin Microbiome Center Gnotobiotics Core Facility. Mice were housed on a 12-h light-dark schedule in a temperature-controlled (22-25°C) and humidity-controlled environment with *ad libitum* access to water and sterile “breeder” chow (Lab Diets 5K52) or experimental diets as described below.

### Gnotobiotic Sequential Feeding Experiment

GF mice at least 6 weeks of age were transferred into flexible film isolators separated by bacterial strain and habituated to the new environment for one week. Mice were then single housed for the duration of the experiment to measure individual diet intake. Mice were introduced to both non-fermentable and fermentable fructan diets (Inotiv-Teklad, **Table S1**)—either inulin and levan diets, or cellulose and inulin diets—for 7 days. On day 8, they were mono-colonized by one 200 µl oral gavage of turbid bacterial culture (∼10^9^ CFU/ml). Mice were then exposed to one of the two diets at a time in sequence, and diet order was counterbalanced in each cohort. Mice were exposed to the first diet for 11 days and to the second diet for 10 days. Throughout these 21 days, daily diet intake was measured every other day, mice were weighed once a week, and feces were collected and plated on BHIS supplemented with 100 µg/ml gentamicin to determine bacterial load on post-colonization days 5 and 15. On day 30, mice were presented both diets at the same time within the home cage one hour before the dark cycle for 16 hours, and overnight intake of each diet was measured. Each sequential feeding experiment was performed two or three times independently, and data were aggregated.

### cFos Immunohistochemistry

Behavior stimuli: GF mice that had gone through the sequential feeding experiment were subjected to both diets on day 32 to trigger acute food preference as a stimulus. Mice were fasted overnight and then habituated to a sterile open field arena for 10 minutes before exposure to both diets for 20 minutes. Following a one-hour rest period for cFos induction, animals were sacrificed by isoflurane, and tissues were fixed via intracardial perfusion of ice-cold PBS followed by 4% paraformaldehyde (PFA). Brains were then post-fixed in 4% PFA at 4°C for 16 hours followed by incubation in 30% sucrose at 4°C for cryoprotection. Tissues were then sectioned at 20 µm and mounted on microscope slides. cFos immunohistochemistry (IHC): IHC was performed as described previously^43^. Briefly, slides were permeabilized in 0.5% Triton/0.05% tween-20 in PBS (PBS-TT) and blocked with 5% normal goat serum in PBS-TT at room temperature for two hours. The tissue sections were then incubated overnight at 4°C with primary antibodies (rabbit anti-cFos (Cell Signaling Technologies) 1:500, and guinea pig anti-NeuN (Sigma-Aldrich), 1:500) in blocking solution (5% NGS + PBS-TT). The following day, slides were washed with PBS-TT before incubating with secondary antibodies (goat anti-rabbit 488 (ThermoFisher) 1:1000, goat anti-guinea pig 568 (ThermoFisher) 1:1000, and DAPI (ThermoFisher) 1:1000) in blocking solution at room temperature in the dark for two hours. Confocal Imaging and Quantification Analysis: Images were obtained and analyzed as described previously^43^. Briefly, three technical replicates of ARH brain tissue were imaged using a 20x air objective (NA 0.8) on an upright Zeiss LSM 780 confocal microscope. ARH sections were selected and ROI’s were drawn based off of the Allen Mouse Brain Atlas. All cell counts were performed by a blinded researcher. First, NeuN positive (NeuN+) cells that were each confirmed to colocalize with a DAPI nucleus were counted. Subsequently, cFos+ cells were counted by confirming colocalization of a cFos immunofluorescence signal with NeuN and DAPI. For each image, the total number of cFos+ neurons were divided by the total number of NeuN+ cells to obtain the percentage of cFos+ neurons. Finally, the percentage of cFos+ neurons for all technical replicates of NTS slices per animal were averaged to obtain a biological n=1.

### Quantification and Statistical Analysis

Statistical analysis was performed using Prism software version 10.1.0 (GraphPad). Data were assessed for normal distribution and plotted in the figures as mean ± SEM. For each figure legend, n = the number of independent biological replicates. Differences between two treatment groups were assessed using two-tailed Student t-test, paired or unpaired as applicable. Differences among >2 groups with 2 variables were assessed using two-way ANOVA and matching across rows with Sidak’s corrections. Significant differences emerging from the above tests are indicated in the figures by #p < 0.10, *p < 0.05, **p < 0.01, ***p < 0.001, ****p < 0.0001. PCA plots were generated in R using the ggplot2 package (v3.4.4)^44^.

## Notes

### Competing Interest Statement

The authors have declared no competing interest.

